# A single-nucleus census of immune and non-immune cell types for the major immune organ systems of chicken

**DOI:** 10.1101/2024.08.05.606414

**Authors:** Elaina R. Sculley, Edward S. Ricemeyer, Rachel C. Carroll, John Driver, Jacqueline Smith, Jim Kaufman, Cari Hearn, Adam Balic, Paula Chen, Susan Lamont, Skyler Kramer, Yvonne Drechsler, Hans Cheng, Wesley C. Warren

## Abstract

In the avian host, comprehensively cataloging immune cell types, their transcriptome profiles, and varying molecular responses to pathogen challenges are necessary steps toward a better understanding of the interplay between genetics and disease resilience. We present a first nuclei atlas of immune cell types derived from the three main immune organs of layer chickens, including spleen, bursa, and thymus. In bursa we also present, an accounting of cell type activation with the bacterial toxin lipopolysaccharide (LPS). Our analysis includes 36,370 total nuclei and 16, 12, and 12 transcriptionally distinct clusters for spleen, bursa, and thymus, respectively. We discover nuclei molecular profiles that uniquely distinguish states of the transcriptome within cell type that could serve as new means to characterize avian immune subtypes. We further subcluster refined immune cell type classifications, specifically highlighting the transcriptomic diversity of B and T cell subtypes. In the bursa, inferred intercellular communication and signaling pathway enrichment analyses across immune and non-immune cell types demonstrate the unappreciated complexity of the B cell repertoire in a model mimicking systemic bacterial infection. This census of all cell types in both primary and one major secondary avian immune organ system, although preliminary, provides a first review of how nuclei transcribe numerous genes, known and unknown, a critical prerequisite for the study avian immunogenetics by cell type.

## Introduction

Chickens serve as invaluable models for studying disease resistance across multiple environments due to their widespread use in both commercial breeding and basic research. With billions of chickens hatched each year, they are among the world’s most important food production animals, and should their resilience to pathogen infection wane, food insecurity would be of immense concern as recently illustrated with influenza A virus outbreaks ^1^. Managing microbial infections in the poultry industry continues to be of high economic value and concern worldwide. Historically, healthy flock management relied on the success of antimicrobial treatment ^2^. However, with increasing pressure to reduce or eliminate antibiotic usage in livestock production, more emphasis has been recently directed toward enhancing host resistance. Gene regulatory responses in the avian host to environmental perturbations are frequently attributed to a diverse range of pathogenic organisms, with their coordinated success determining animal well-being.

To counteract the diversity of pathogens, vertebrates have evolved a complex interplay of innate and adaptive immunological mechanisms to enhance their chances of survival. The complement system, functioning as a cascade, liberates antimicrobial components and factors that attract and modulate the cellular immune system ^3^. Innate responses play a crucial role in the early stages of microbial invasion, limiting pathogen spread until the adaptive immune response is activated which involves highly specific T- and B cell-mediated mechanisms to eliminate infections. Unlike the thymus which produce naïve B and T cells, respectively, the immune cells of the spleen compositionally enable an efficient mounting of both innate and adaptive immune responses. Therefore, a comprehensive study of all three tissue types is warranted to better understand the molecular features of strong immune resistance for future genetic selection strategies ^4^.

In large-scale layer and broiler producing environments with high bird-to-bird contact, chronic exposures to many evolving pathogens continue to be problematic for the poultry industry. In birds with lower tolerance or in stressful states, costly loss of productivity occurs. Natural antibodies to foreign antigens exist in all birds and have been linked to better survival in chickens ^5^ ^6^ but the practicality in selective breeding for these antibody responses remains unproven. Previous chicken studies have revealed a complex network of immune responses initiated by various bacterial pathogens ^7^. Increasing interest in the sophistication of host immune response has enhanced research efforts to decipher the underlying cellular mechanisms, yet a vast majority of our avian host response knowledge is based on cell sorting, tissue histology, and bulk RNAseq data without a comprehensive representation of all cell types within the tissue ^8^. For the resistance phenotype, there is genetic evidence for some line-specific consistency to two of the most significant diseases affecting poultry, coccidiosis and necrotic enteritis, nonetheless the authors concluded informed breeding is still not possible ^9^. A significant locus of ∼1Mb harboring two candidate genes was identified suggesting involvement in Salmonella resistance, but again, an accounting of their collective molecular roles was missing ^10^.

One means to effectively study *in vivo* bacterial infection dynamics is exposure to LPS, a component of the cell wall of Gram-negative bacteria, LPS, a representative pathogen-associated molecular entity to which the immune system mounts an immediate response ^11^. LPS in a stepwise way interacts with a protein complex of the CD14/TLR4/MD2 receptor complex present in host immune innate and adaptive cell types, e.g. macrophages, but the subsequent response depends on the cell type ^12^. In the infected bird, dividing bacteria undergo autolysis stimulating immune cell-mediated killing that occurs via complement activation, phagocytosis, or by B cell-produced antibodies. Interestingly, different LPS entities can illicit low or high immunogenic responses for some bacterial species. A low immunogenicity version of LPS is suggested to be a strategy to evade host immune response and increase intracellular survival, the so-called good bacteria ^13^. Although much has been learned about how the chicken responds to bacterial infections, both in terms of specific pathogens as well as broader phyla representation, it is now essential to build a cell type-specific contextual framework for understanding how systemically pathogens can overcome the host immune response.

Single-cell (scRNAseq) or single-nuclei RNAseq (snRNAseq) approaches have fundamentally changed how we now go about reconstruction of distinct cell populations and their gene expression patterns ^14^. Large consortium projects are providing new vital landscapes, immune and non-immune cell types, predominately for mouse and human ^15^ ^14,16^. Cell type-specific gene expression profiles from varying organ systems at large scale 360,000 cells, some non-immune centric, demonstrated distinct and shared features for various immune cell types, e.g. macrophages ^16^. Using this approach, these analyses discovered unappreciated tissue-specific features of immune cell systems, such as the full diversity of the lymph node that is now nine peripheral lymph node non-endothelial cell clusters, thus suggesting new niche-restricted immune functions ^17^. Very few studies have evaluated the differences in single cell and nuclei transcriptomes, but importantly the ability to annotate cell types with both methodologies was mostly equivalent ^18^ ^14^. Wu et al. point out snRNAseq offers compelling advantages to scRNAseq because samples are cryopreserved for experimental flexibility, better representation of the original cellular state compared to live cell protease perturbations, and gene regulatory events that govern cell dynamics are enriched, e.g. transcription factors ^18^. Studies that provide an immune cell-type resolution by organ system with concomitant nuclei gene expression signatures are not yet available in the chicken. Should cell type atlases become available in chicken, albeit at smaller scales, when integrated with the human and mouse cellular data, a new in-depth knowledge of the avian immune system can be constructed, which are thus far, only classified with antibodies to cell surface antigens.

Recently, studies have begun to deploy single cell technology in attempts to decipher avian host immune response by cell type as a start to understand unaccounted sources of variation ^19–24^. Although most have focused on viral infection and the use of live cell preparations, many new findings have been uncovered. From these experiments some examples of ligand-receptor candidates were offered for the viral pathogenesis of Newcastle Disease infection ^21^, the evolutionary conserved role for XCR1+ cDCs in monitoring CD8+ T-cells biology ^22^, and Marek’s viral disease in the spleen that show some cell types are more vulnerable ^23^.

Because genomic-assisted selection programs continually attempt to build disease resilient flocks, it is essential that the networks of immune genes intricately associated with myeloid and lymphoid cell types reach higher levels of resolution so their importance can be better measured. We contend cell types identified by their single nuclei or cell transcriptome profiles will allow for their expanded integration and interpretation. In this study, we used a well-characterized Avian Disease and Oncology Laboratory (ADOL) White Leghorn line to generate a first census of immune cell types in the major immune organs of chicken, the spleen, bursa, and thymus (Fig. 1). From these cell-specific results, we provide a novel single nuclei transcriptome perspective and postulate various systems biological interpretations of the chicken’s bursal immune response to bacterial infection.

**Figure 1.**
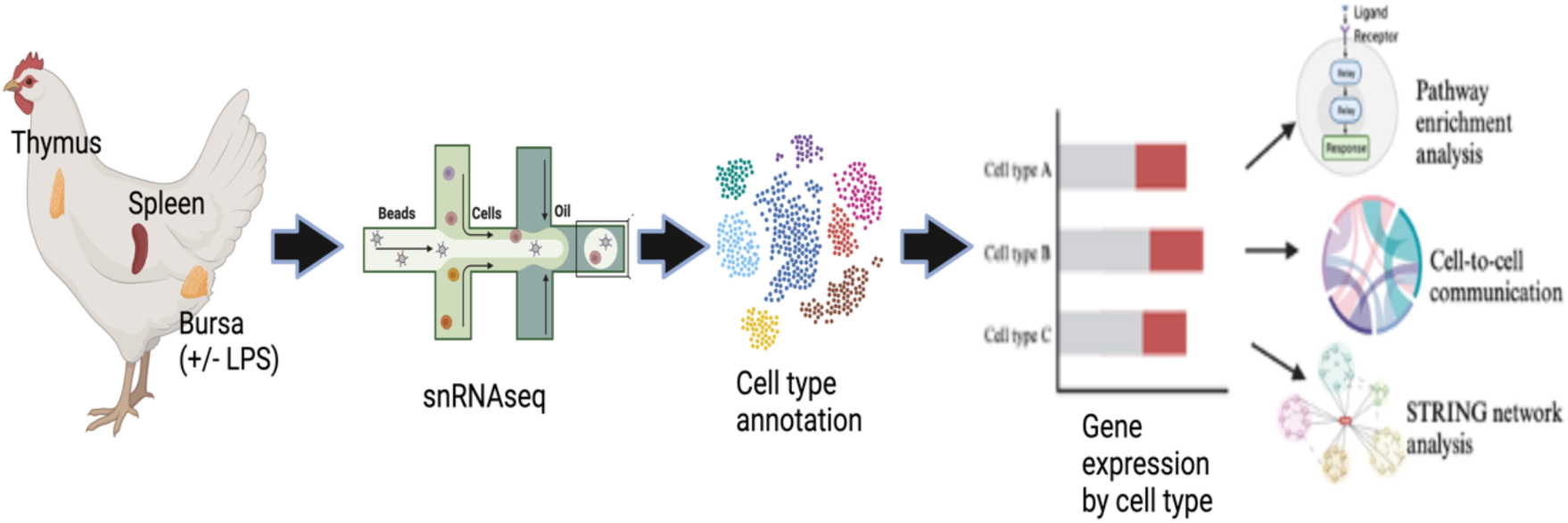
A schematic of objectives and the experimental design to characterize immune cell types in the chicken. Image created with BioRender.com.

## Materials and Methods

### Sample processing

We used Avian Disease and Oncology Laboratory (ADOL) female White Leghorn line 6 birds ^25^ and performed snRNAseq to determine the cell-type transcriptomes of the bursa, spleen, and thymus plus transcriptomic changes post-LPS administration for the spleen and bursa only (Fig. 1). Birds at five weeks of age were injected intravenously with a dose of 1 mg/kg LPS-derived from *E. coli* (strain 055:B5, catalog# L2880 Millipore Sigma), and biphasic body temperature response to systemic LPS treatment was measured at 0, 1, 4, 10, and 24h (Suppl. Table 1). Samples for snRNAseq data generation were collected at between 6.75-9.75 hours post-LPS injection. For these experiments, we used CO_2_ gas euthanasia, following the current standards for poultry euthanasia provided in AVMA Guidelines for the Euthanasia of Animals (2020 Edition). All studies presented herein were carried out in accordance with the approval of the Institutional Animal Care and Use Committee, USDA, ARS, ADOL, East Lansing, MI (protocol approval number 2018-01). Moreover, all methods were performed in accordance with the ARRIVE guidelines ^26^. Upon dissection, each tissue was flash frozen in liquid nitrogen for long term storage at -80°C.

### Nuclei isolation and sequencing

Our decision to use nuclei instead of cells was based on the use of flash frozen samples that permits measures of the transcriptome not provoked by stress during mechanical and protease treatments, future parallel studies of externally collected and cryopreserved samples, the broadly comparable detection of the same cell types, and an ability to accurately annotate immune cell types as demonstrated in Eraslan et al. ^14^.

Nuclei were isolated from spleen and bursa as previously described in Eraslan et al. ^14^, while for thymus, we used the Singulator2 instrument per manufacturer’s protocols (S2 Genomics; Livermore, CA). After mechanical disruption, cell filter straining steps, and washes, the cell nuclei were suspended in PBS/0.1% BSA or for thymus a nuclei isolation buffer (S2 Genomics; Livermore, CA), with all buffers containing 0.4U/ul RNase inhibitor (source), prior to microfluidic encapsulation on the 10X Chromium instrument (10x Genomics®, Pleasanton, CA) to nanoliter-scale Gel bead-in-EMulsions (GEMs). Single-nuclei libraries were generated using the GemCode Single-Cell Instrument and Single Cell 3′ Library and Gel Bead Kit v3 and Chip Kit (10x Genomics®, Pleasanton, CA) according to the manufacturer’s protocol. Before sequencing, every library was analyzed on a Bioanalyzer high sensitivity chip to ensure the expected cDNA fragment size distribution was achieved. The appropriate number of individually barcoded GEM libraries were pooled and sequenced on a NovaSeq 6000 instrument (Illumina) with 2×150bp length using these sequencing parameters: 26 bp read 1 – 8 bp index 1 (i7) – 98 bp read 2 with 200 cycles.

### Single-nuclei RNAseq preprocessing

The Cellranger software pipeline (version 3.1.0) was used to demultiplex cellular barcodes, map reads to the GRCg7b chicken reference genome ^27^ and transcriptome using the STAR aligner, and down-sample reads required to generate normalized aggregate data across samples, producing a matrix of gene counts versus cells. The individual tissue-specific sequenced Gel Bead-In Emulsion (GEM) libraries were each processed with the Cellranger v7.0.1 pipeline (10X Genomics) to create a cellular barcode by genomic feature matrix that was then analyzed using the Scanpy tools v1.9.2 ^28^. To remove detectable ambient RNA, CellBender ^29^ was used on all raw gene expression matrices using the remove-background-v2-alpha workflow with FPR=0.01 option. In preparation for clustering, we filtered out nuclei using the distribution of detected genes, mitochondrial and ribosomal transcripts specific to each tissue source. We normalized and log-scaled the counts matrix, computed highly variable genes, regressed out counts per cell and performed principal component analysis (PCA) prior to integration across samples using harmonypy v0.0.9 ^30^. We computed the 10-neighbor graph with the first 20 adjusted principal components clustering using the Leiden algorithm ^31^ with resolution 0.5, and visualized in two dimensions with uniform manifold approximation and projection (UMAP) ^32^. A Jupyter notebook containing all code used in this analysis is available along with this paper.

### Cell type annotation

We produced a list of marker genes for each cluster by performing a t-test with overestimated variance to find genes with significantly higher expression in the cluster in question compared to all other clusters (LFC > 1; p < 0.01) (Wolf, Angerer et al. 2018)). Our nuclei dataset is novel for the chicken, essentially, we do not have nuclei-specific markers for any cell types. Therefore, we implemented a manual process that used a combinatorial approach, including known cell-type chicken specific gene markers (e.g. *PAX5* for B-cells), team curation by avian immunologists each evaluating the significant differentially expressed genes (DEGs) per cluster (p value<0.01), and comparisons to existing mouse and human scRNAseq databases of CellMarker ^33^, ImmGen ^34^, and PanglaoDB ^35^.

### Cell type composition and gene expression comparisons

After cell type identities were finalized, we plotted their composition by organ source and for LPS treatment effects, spleen and bursa, then estimated any significant changes using scCODA ^36^. For non-LPS controls of thymus, we simply counted nuclei belonging to each cell type and plotted the percent contribution of each.

### Differential gene expression by treatment

To compute genes that were differentially expressed in each cluster between the control samples and the LPS-treated samples, we summed raw counts for all cells in each sample to obtain a pseudo bulk counts matrix, as this method has been shown to perform better than those that treat each cell in a sample as an independent data point ^37^. We then used this pseudo bulk counts matrix as input to pyDESeq2 v0.3.5 ^38^, which we ran with the default parameters. We used a minimum absolute value of log2FC of 1.0 and maximum adjusted p-value of 0.01 for a gene to be considered differentially expressed. The only adjustable parameter the method considered was a threshold for the minimum percentage of cells in each cluster that express a gene for it to be considered; we used the default cutoff of 0.1.

### Cell to cell communication analysis

Cell-cell interactions in the bursa for all cell types were analyzed using CellChat v2 ^39^. We converted chicken gene names using previously defined gene orthology to the published human dataset of ligand-receptor pairs in the CellChat database to test for enrichment of ligand-receptor interactions within and between cell types. Interaction heat maps were built using the outgoing or incoming signals as curated in the ligand-receptor pairs database as a module of CellChat. The labeled intensity of coloration per gene is in accordance with the observed level of ligand or receptor transcript counts in each cell type (p value threshold of 0.05).

### Signaling pathway enrichment in bursa

To evaluate enrichment across all cell types in bursa, we calculated the hypergeometric distribution of overlapping genes over all genes in a given collated test set as defined in the Molecular Signatures Database’s of canonical pathways (v7.2) ^40^. A statistical test was performed on a hypergeometric distribution of overlapping genes over all genes in the universe (annotated collection) to calculate p-values. We report the false discovery rate (FDR) analog of the p value after correction for multiple hypothesis testing as the FDR q-value for each pathway above the p-value of 0.05. We were guided by the methods of Espinoza et al. ^41^ to characterize B cell pathway enrichment.

### String network analysis

The DEGs were used as input for bursa and investigated by overlap with known protein-protein interactive networks using STRING ^42^ as described previously ^23^.

## Results

### Initial clustering of all bursa cell types

For all samples (2 LPS and 3 control), a total of 14,923 nuclei were partitioned to form 16 transcriptionally distinct clusters. Manual inspection of the cluster plots distribution of individual cells were mostly uniform between LPS and control groups for bursa and spleen (Suppl. Fig 1A-D). A mean of 1,153 genes were detected per nucleus (Suppl. Table 2). The manual cell type identification of each cluster relied on a convergence of multiple databases defining markers associated with specific cell types (Fig. 2A; Suppl. Fig. 2). Multiple examples of bursa gene markers displaying their expression distribution across individual nuclei and clusters are provided in Suppl. Fig. 2.

**Figure 2.**
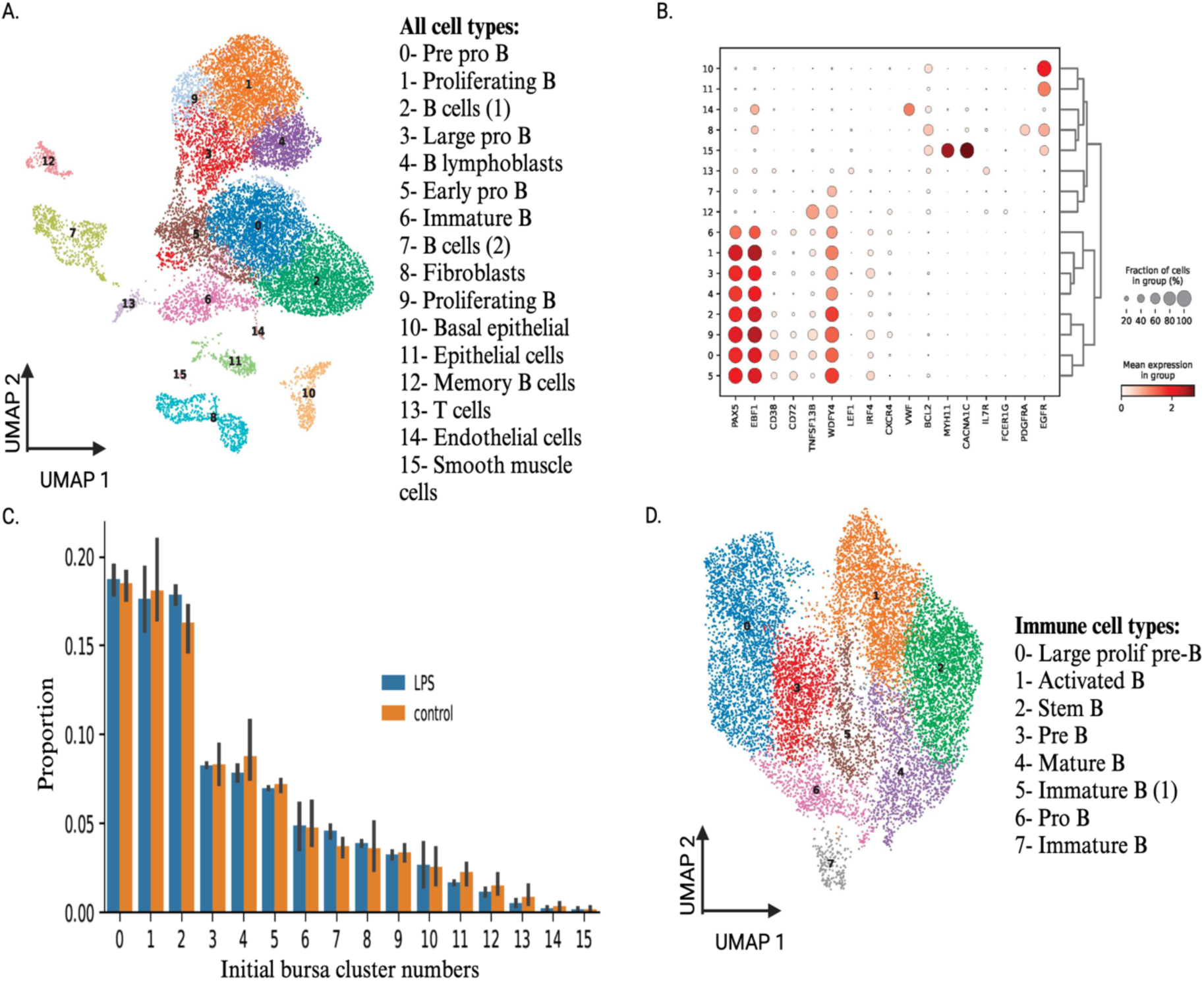
Single-nuclei transcriptomic analysis of the chicken bursa. **(A)** Uniform manifold approximation and projection (UMAP) visualization of bursa cell types. Clusters were identified using the graph-based Louvain algorithm at a resolution of 0.5. **(B)** Dot plot showing the Z-scored mean expression of marker genes used to designate cell types to cell clusters. The color intensity indicates the average expression of each marker gene in each designated cell type. The dot size represents the proportion of cells which expression of the gene was detected. All genes with significant expression (p<0.01) that defines a cluster relative to all other clusters are presented in Suppl. Table 3. **(C)** Composition differences by cell subtypes. **(D)** Subclustering of all identified B cells in the initial clustering and their annotation. Image created with BioRender.com.

### Bursa lymphoid

Most clusters (0-7, 9, and 12) are identified as B cells using common B cells markers, *CD38*, *PAX5, EBF1*, and *CD72* (Fig. 2B; Suppl. Fig. 2). Variable expression of these B cell markers suggested different B cell subtypes or varied states of expression for major B cell types. We attempted to assign B cell developmental stages to each cluster using the stages previously described ^43^. Cluster 0 is named pre pro B based on expression of *TNFSF13B*, which codes for the BAFF receptor that binds the B-cell activating factor (*BAFF*), a regulator of the peripheral B-cell population. Clusters 1 and 9 are predicted to be proliferating B cells based on the dominant expression of *POLA1* (Suppl. Fig. 2; Suppl. Table 3). Cluster 2 displays a similar B cell marker expression pattern with other B cell clusters (Suppl. Table 3) as well as some not well-characterized genes, such as the *WDFY4* expressed gene that is associated with B cell survival during autophagy ^44^. In clusters 3 and 5, expression of *JCHAIN*, a protein responsible for binding activity in dimers of Immunoglobulin M and Immunoglobulin A (Frutiger et al. 1992), suggests a plasma B cell phenotype (Suppl. Table 3). Moreover, the *IRF4*, which encodes for a protein involved in IgA class switching role, is expressed in these clusters (Suppl. Table 3). In cluster 7 there is a small amount of B cell marker-high cells but the important myeloid marker genes as well (*MARCO, MAFB, TIMD4, FLT3*) suggest these cells could be B cell-efferocytosing macrophages (Suppl. Table 3). For cluster 4, co-expression of *RAG2* with *POLA1*, and *BRCA1* suggest a B lymphoblast subtype. Cluster 12 is memory B cells based on *PRDM1* and *CD86* gene expression. A small population of T cells, cluster 13, are positive for *TCF7* (Suppl. Fig. 2), *BCL11B,* and *TARP* (Suppl. Table 3).

### Bursa non-immune cell types

In order of their observed overall abundance were epithelial (clusters 10 and 11), fibroblasts (cluster 8), endothelial (cluster 14), and smooth muscle cells (cluster 15). Distinguishing gene markers for each are shown in Fig. 2B. Epithelial cells gene expression of *TP63* and *PPL* are one example (Suppl. Fig. 2).

### Bursa cell composition of all cell types

We estimated the proportional changes for all cell types, immune and non-immune, using scCODA ^45^. No cell types showed a significant shift in proportion as a result of LPS treatment (Fig. 2C).

### Bursa subclustering of B cell types

To further characterize subtypes, we subclustered nuclei previously identified as B cells (n=12,608) that resulted in nine transcriptomic distinct clusters (Fig. 2D). Of the total bursa nuclei collected, this shows that 84% are B cells of which we display clusters 0-7 since 8 was of unknown origin (Fig. 2D). Most stages of B cell development, including pro-B, pre-B, and immature, and mature B cells, were present according to gene markers that define B cell trajectories in the mouse ^43^. The expression of *EBF1*, a pan B cell marker, shows the subclustering success of identifying only B cells from all cell types (Suppl. Fig. 3). Clusters 0 and 3 represent large pre-B cells in different states (Suppl. Fig. 3; Suppl. Table 3). While cluster 0 expressed genes, including *RAG2*, *VPREB*3, and *CD1C,* suggestive of pre-B status, its coexpression of the proliferation marker *POLA1* also supports a rapidly dividing molecular phenotype (Suppl. Fig. 3; Suppl. Table 3). Cluster 1 is possibly an activated B cell based on *TNFSF10* and *CLEC19A* expression (Suppl. Fig. 3; Suppl. Table 3). Cluster 2 appears to be stem B cells as evidenced by *CD38, ATXN1,* and *SOX4* gene expression (Suppl. Fig. 3; Suppl. Table 3). Cluster 4 is mature B based on the expression of *CD72, BCL2, and CD74,* the latter playing a cell differentiation role (Suppl. Fig. 3; Suppl. Table 3). Cluster 5 is annotated as immature B using *EIFEBP1* and *CXCR4* (Suppl. Table 3). Cluster 6 is pro-B relying on *IGLL1*, *JCHAIN,* and *IKZF1* expression signatures (Suppl. Table 3). Expression of the *IKZF1* gene, a transcription factor, is critical for pro-B cell commitment. Cluster 7 is immature B with *RUNX2, CD247*, and *UTRN* (Suppl. Table 3).

## Spleen

### Spleen initial clustering of all cell types

The spleen is capable of both innate and adaptive immune responses and is highly compartmentalized with each displaying its own arrangement of lymphoid inducer tissues in T and B cell-dependent areas ^46^. For five individual samples, two control and three LPS treated, we obtained a total of 10,233 nuclei that passed our quality filtering for further analysis. A mean of 687 genes showed detectable expression per nucleus. Other quality metrics are in Suppl. Table 2. The Leiden clustering of all nuclei led to 12 transcriptionally unique clusters with the largest being a population of T cells (Fig. 3A). Of the 12 clusters, four were determined to be the non-immune cell types, endothelial and fibroblasts, each displaying different transcriptomic states (Suppl. Fig. 4; Suppl. Table 3).

**Figure 3.**
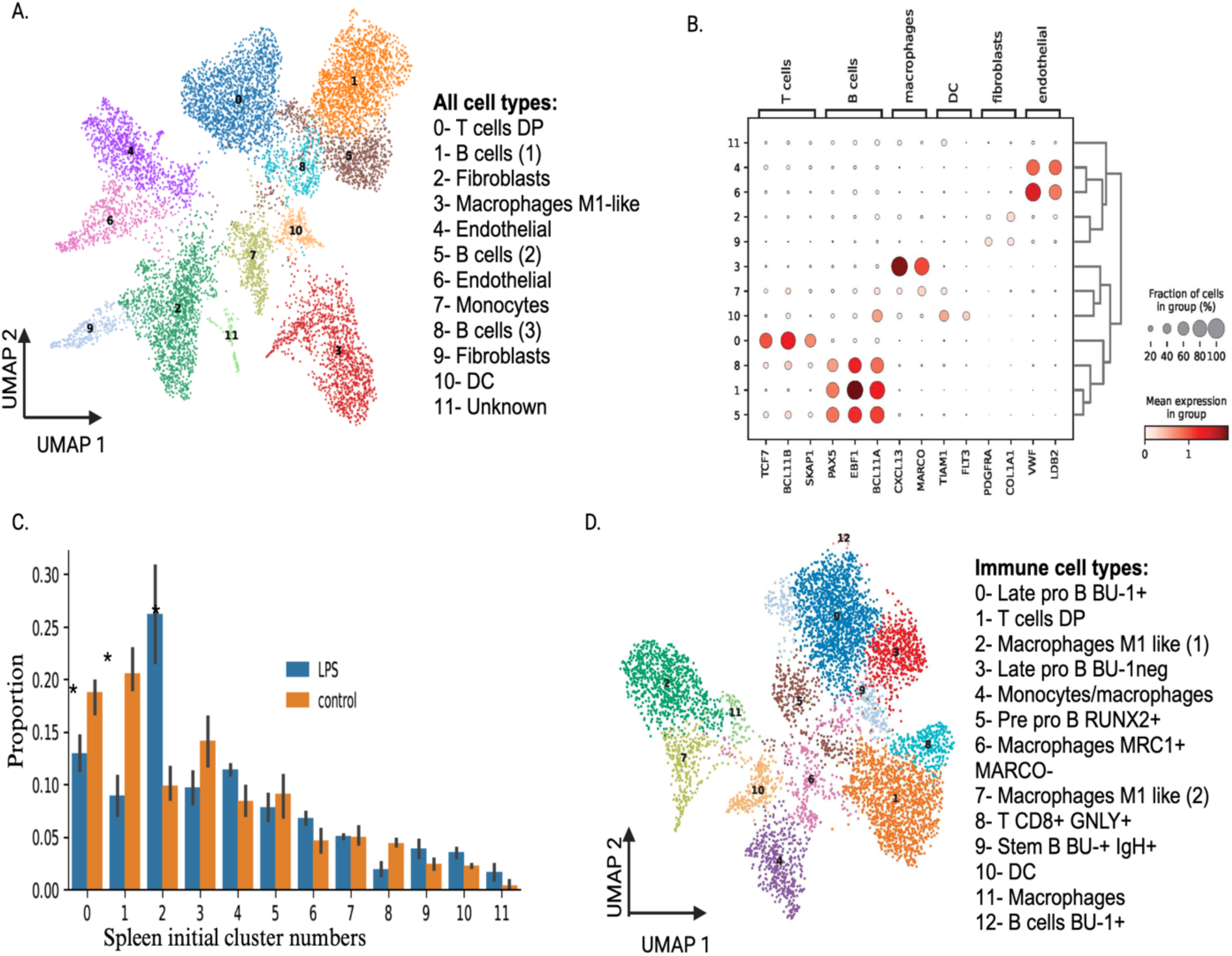
Single-nuclei transcriptomic analysis of the cellular composition of the chicken spleen. **(A)** Uniform manifold approximation and projection (UMAP) visualization of spleen cell types. Clusters were identified using the graph-based Louvain algorithm at a resolution of 0.5. **(B)** Dot plot showing the Z-scored mean expression of marker genes used to designate cell types to cell clusters. The color intensity indicates the average scaled expression of each marker gene in each designated cell type. The dot size represents the proportion of cells for which expression of the gene was detected. All genes with significant expression (p<0.01) that defines a cluster relative to all other clusters are presented in Suppl. Table 3. **(C)** Composition differences by cell subtypes. **(D)** Subclustering of all identified lymphoid and myeloid immune cells in the initial clustering and their annotation. Image created with BioRender.com.

### Spleen lymphoid

The largest and only T cell population, cluster 0, is enriched with DP T cells (*CD4, CD8A;* Suppl. Table 3) but also contains cells expressing classic T cell markers including: *BCL11B, TCF7, CD3E, SKAP1,* and *TARP* (Fig. 3B; Suppl. Fig. 4). Further manual review suggests smaller populations of single positive *CD4+* and *CD8A+* cells are present as well and clustered together with DP T cells. For B cells, we identify three clusters 1, 5, and 8 showing coexpression of *PAX5, EBF1,* and *BCL11A* genes, indicating this is proportionally the dominant cell type in spleen (Fig. 3B; Suppl. Fig 4). Differences are also observed, such as higher expression of *CD38* in cluster 1 (Suppl. Fig. 4) and cluster 8 displaying a proliferating B cell signature based on *BRCA1, CENPF,* and *KIFC1* expression (Suppl. Table 3).

### Spleen myeloid

Three clusters, 3, 7, and 10, were associated with the innate immune system. Cluster 3 is macrophages based on defining gene markers, *CXCL13* and *MARCO (*Macrophage receptor with collagenous structure), with an M1-like phenotype due to *CD86* coexpression (Fig. 3B; Suppl. Fig. 4). Cluster 7 appears to be a small grouping of monocytes (*KUL01* and *TIMD4*), and cluster 10 is dendritic cells (DC) as shown with *FLT3, TIAM1, CADM1,* and *IRF8* gene expression (Suppl. Fig. 4; Suppl. Table 3).

### Spleen non-immune cell types

Clusters 2, 4, 6, and 9 were identified as various non-immune cell types (Fig. 3B; Suppl. Fig. 4). These in order of proportion were endothelial cells (clusters 4 and 6) and fibroblasts (clusters 2 and 9), each with defining gene markers (Suppl. Fig. 4). For example, *VWF* gene expression marking endothelial cells (Fig. 3B; Suppl. Fig. 4).

### Spleen composition for all cell types

Upon prediction of cell types, we estimated the proportional changes induced by LPS for each cell type using scCODA, a Bayesian method described in ^36^. In comparing LPS against control, we find a significant decrease in the proportion of cells associated with adaptive immunity, i.e. number of B and T cells, in LPS-treated birds, and an increase in proportion of macrophages (Fig. 3C), suggesting a shift from adaptive to innate immunity upon LPS treatment.

### Spleen subclustering of only immune cell types

All nuclei associated with identified immune cell types (n=6,804) were subclustered, resulting in a total of 13 distinct clusters (Fig. 3D). Of all spleen cell types, 66% are of immune origins in this study. One large T cell population previously enriched for *CD4* and *CD8A* markers (Suppl. Table 3) separated into two clusters, one a mixture of single positive *CD4* and *CD8A* expressing cells and the other classic cytotoxic T cells, the latter distinguished by *CD8A* and *GNLY* expression (Fig. 3D; Suppl. Fig. 5). For the innate system, one initial macrophage population (Fig. 3A) was subsequently resolved into four subtypes, clusters 2, 6, 7, and 11, with varied scavenger receptor *MARCO* expression as a distinguishing feature (Fig. 3D; Suppl. Fig. 5). All gene markers used to define subclustered immune cell types of the spleen are summarized in Suppl. Table 3.

## Thymus

### Thymus initial clustering of all cell types

After quality filtering, two samples yielded a total of 11,214 nuclei (Fig. 4A). A mean of 561 genes per nucleus showed detectable expression. DEGs per cluster that define cell type are available in Suppl. Table 3. Other metrics of quality are available as well (Suppl. Table 3). The initial Leiden clustering of all nuclei led to 12 transcriptionally unique clusters with the largest being a population of double positive (*CD4* and *CD8A*) T cells (Fig. 4A). Only clusters 8 and 10 were identified as non-immune, epithelial and erythrocytes (Fig. 4A; Suppl. Fig. 6). Erythrocytes (cluster 10) were removed from Fig. 4A.

**Figure 4.**
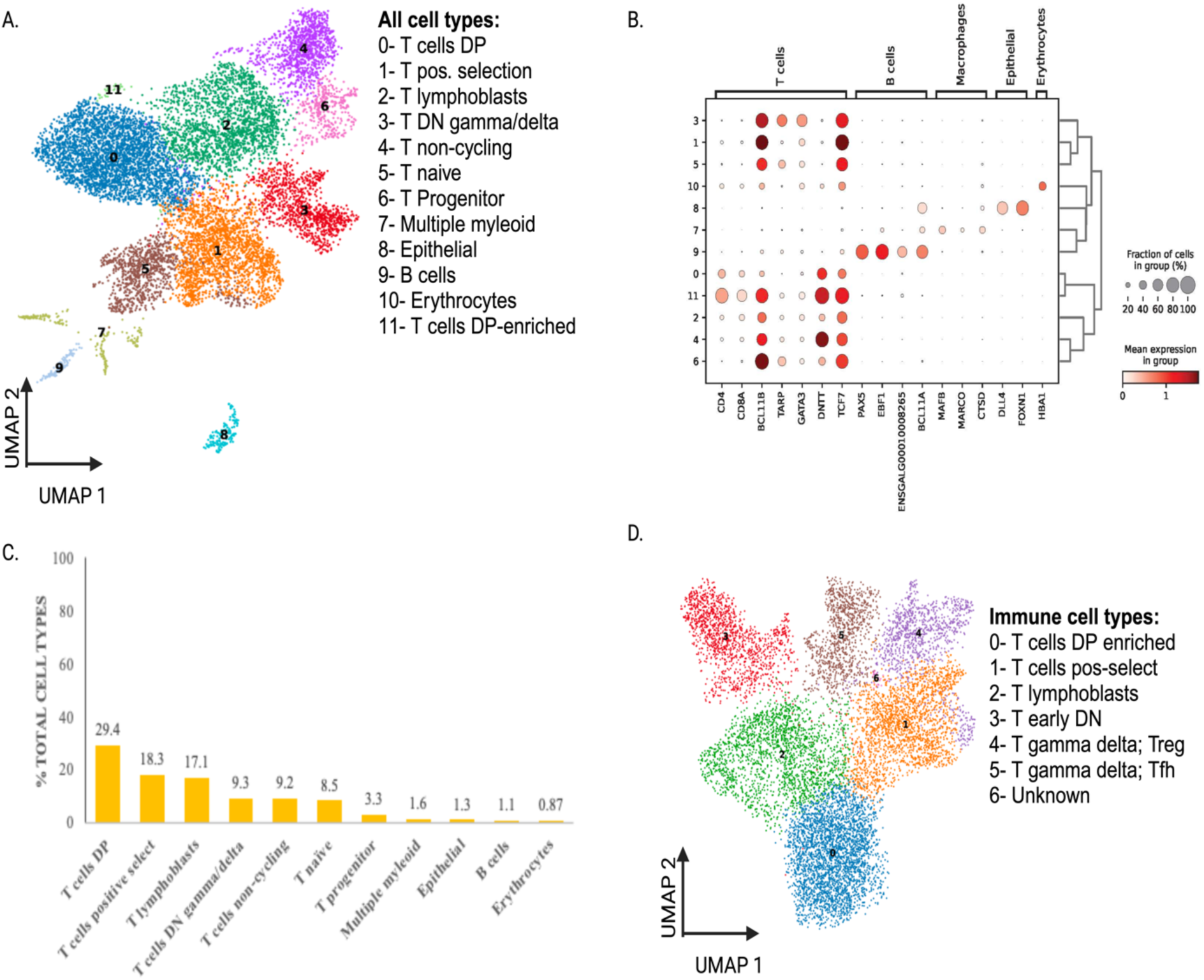
Single-nuclei transcriptomic analysis of the cellular composition of the chicken thymus. Uniform manifold approximation and projection (UMAP) visualization of thymus cell types. Clusters were identified using the graph-based Louvain algorithm at a resolution of 0.5. (B) Dot plot showing the Z-scored mean expression of marker genes used to designate cell types to cell clusters. The color intensity indicates the average scaled expression of each marker gene in each designated cell type. The dot size represents the proportion of cells for which expression of the gene was detected. All genes with significant expression (p<0.01) that defines a cluster relative to all other clusters are presented in Suppl. Table 3. (C) Composition differences by cell subtypes. (D) Subclustering of all identified T cells in the initial clustering and their annotation. Image created with BioRender.com.

### Thymus lymphoid

Most clusters (0-6 and 11) are identified as various types of T cells using well-known T cells markers: *TCF7, CD8A*, and *CD4* (Fig. 4B). Cluster 2 represents a proliferating state with the expression of *POLA1* and *CENPF* (Suppl. Fig. 6). Three clusters (3, 5, and 6) suggest gamma delta T cell lineages based on higher *TARP* expression (Fig. 4B). However, Cluster 6 seems unique as a grouping of double negative cells with some early/cortical/DN pre-T-consistent markers (*CDK6, KIT, CD44, PTPRK*), as well as a proliferation signature (Suppl. Table 3). Clusters 4 and 11 appear to be early developing T cells based on *DNTT* expression that is associated with T cell hierarchy, but cluster 0 also shows moderate expression (Suppl. Table 3). For other adaptive cell types, we identified a small population of B cells, cluster 9, that express *PAX5*, *EBF1*, ENSGAL00010008265 (*BU-1*), and *BCL11A* (Suppl. Fig. 4B).

### Myeloid

Only a fragmented cluster 7 was of myeloid origins, which we label as macrophages based on *MAFB*, *MARCO,* and *CTSD* expression patterns (Fig. 4B). One fragment of this cluster is likely DC due to the expression of *FLT3* (Suppl. Fig. 6).

### Thymus non-immune cell types

The largest non-immune cell types proportionally were epithelial (cluster 8) with the *NPAS3* gene as the best marker to differentiate this cell type. We predict this cluster to be cortical thymic epithelial-like based on the expressed genes, *DLL4* and *FOXN1* (Fig. 4B). We also see erythrocytes with the prominent expression of *HBA1* (Fig. 4B), *ANK1,* and *HBBA* (Suppl. Fig. 6).

### Subclustering of all T cell types

We identified seven transcriptionally distinct clusters when starting with all nuclei initially identified as T cells (n=10,704) with one of unknown identity (Fig. 4D). In this small sample set, this shows that 95% of cells in the thymus are T cells. Each cluster displayed combinations of gene markers that allow for their unique subtype annotation. However, low mean genes detected per nuclei prevented complete T cell subtype classifications. Cluster 0 is double positive T cells, while cluster 2 is likely T lymphoblasts due to the abundance of the proliferation markers *BRCA1* and *POLA1* as well as T cell-specific genes (Suppl. Fig. 7). Cluster 3 appears to be double negative early progenitor T cells using the *DNTT* and *PLD5* markers as evidence (Suppl. Fig. 7). Cluster 4 is predominantly gamma/delta T cells based on *TARP* as well as some cells expressing T regulatory markers, such as *LAG3* and *KLF2*, the latter is a marker of T cell egress (Suppl. Fig. 7). Cluster 5 is also gamma/delta T cells but with a marker that is affiliated with T follicular helper cells, *BCL6* (Suppl. Fig. 7).

## Immune system activation

### Bursa all cell types’ nuclei transcriptomic response to LPS

Using a pseudobulk approach ^37^ to find genes differentially expressed between control and challenge conditions in each cluster, we find highly variable gene regulation across all cell types (Fig. 5A). The most transcriptionally active compared to all others was the basal epithelial cells (Fig. 5A; Suppl. Table 4). Overall, we find little overlap between the four largest B cell clusters (Suppl. Fig. 8).

**Figure 5.**
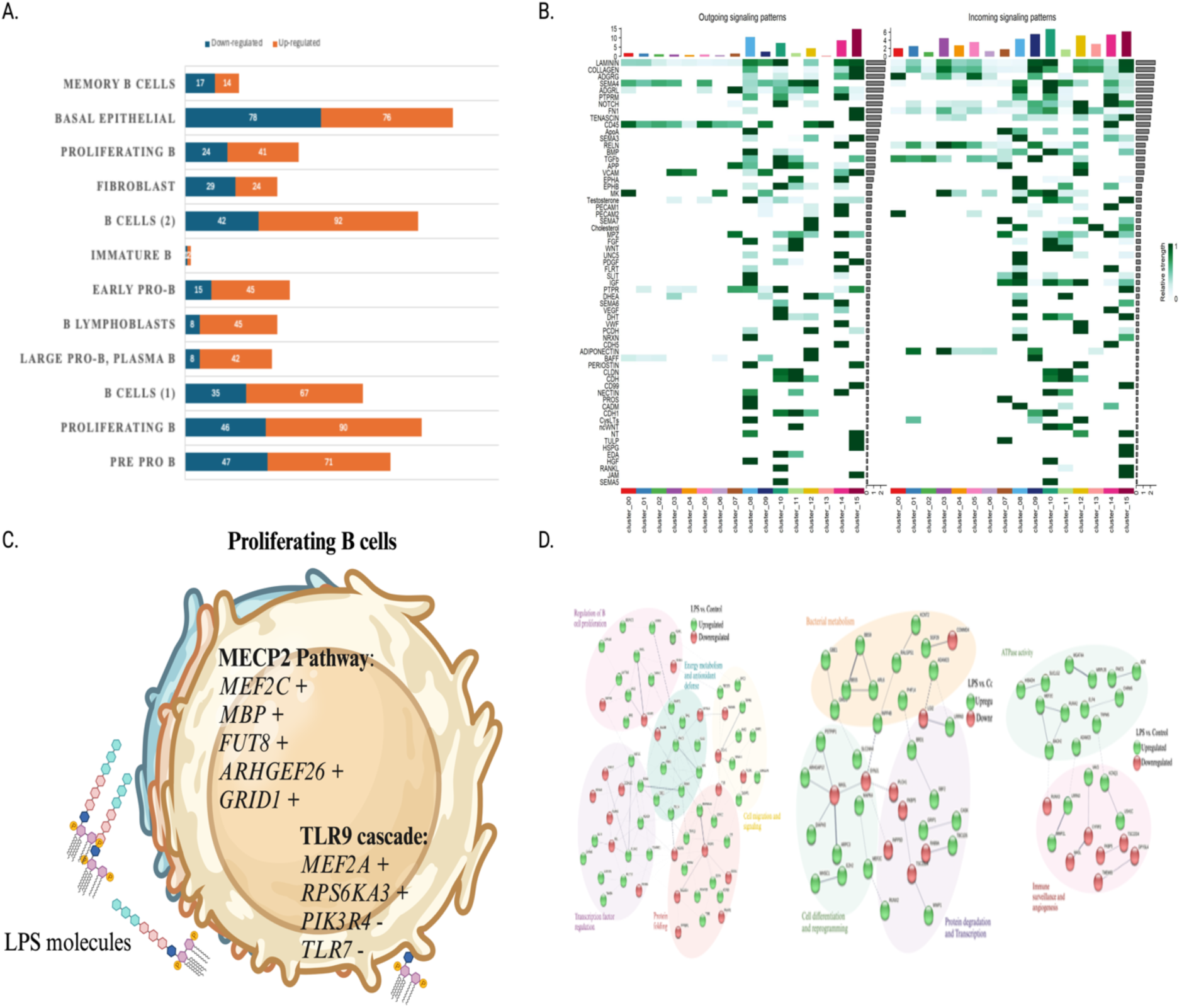
Bacterial toxin activation of bursal cellular communication and signaling. (A) summary of up (orange) and down (blue) DEGs by cell type and (B) analysis of cellular crosstalk with CellChat for all interactions and those meeting significance criteria for incoming and outgoing signals. (C) Canonical pathway enrichment using MySigDB for the proliferating B cells subtype for MECP2 and TLR9 signaling and (D) STRING network analysis of DEGs in the subclustered B cells. Image created with BioRender.com.

### Bursa cell-to-cell communication for all cell types

After performing CellChat across all cell types, we reviewed significant intracellular and intercellular interactions, both outgoing (ligands) and incoming (receptors) between specific cell type pairs within the broader landscape of all cell-to-cell interactions (Fig. 5B). Specific interactions that occurred are highlighted for all interactions, and our analysis revealed significant interactions between specific B cell ligand-receptor pairs in various immune cell interactions (Fig. 5B). We see the interaction between *SEMA4D* (Semaphorin 4D) and *CD72*, especially in the context of B-cells and immune regulation, which are both cell surface proteins where their interaction plays a crucial role in immune cell signaling and modulation ^47^. Additionally, we see the interaction between *PECAM1* and *CD38*. *PECAM1* is a cell adhesion molecule that plays a role in leukocyte trafficking ^48^, and *CD38* is a multifunctional cell surface glycoprotein ^49^; both are expressed on a variety of immune cells including B cells.

### Signaling pathway enrichment for all cell types in bursa

The evaluation of cellular signaling and its enrichment was accomplished using the MySigDB pathway enrichment analysis of the DEGs per cell type in the bursa (Suppl. Table 4). We find enrichment signals because of LPS activation for most cell types, but some (epithelial, T cells, endothelial, smooth muscle cells) did not return DEGs using our statistical criteria (see methods; Suppl. Table 5). For a targeted set of cell types including pre pro, proliferating, and B cells (1) and (2), basal epithelial, and fibroblasts, we report pathway descriptions, genes in overlap, and the FDR q-value scores for each (Suppl. Table 5). When examining these different states of B cell LPS activation, some commonalities emerge among them; pre pro B and proliferating cells for genes associated with the *MECP2* pathway and epithelial TP63 and basal epithelial cells overlapping in the rho GTPase pathway (Suppl. Table 5). The rho GTPases are pleiotropic; involved in many biological processes, including lymphomas indirectly connecting their role to immune response ^50^. Many genes of interest were considered, but we highlight just one perhaps expected finding in proliferating B cells, multiple regulated genes associated with the MECP2 pathway (Fig. 5C and D). *MECP2* functions as a transcriptional repressor and activator ^51^. In addition, we find involvement of the Toll like receptor family, specifically TLR9 pathway, a known contributor to innate immune responses (Fig. 5C).

### Protein-to-protein interaction

To gain insight into the DEGs that are expressed by the subclustered cell types of B cells, we ran STRING network analysis ^42^. By serving as a meta-database, STRING aligns interaction evidence across various genomes and proteins, encompassing approximately 2.5 million proteins from 630 organisms ^42^.

When searching protein-protein interaction networks using on B cell DEGs, we find multiple genes associated with immune functions such as positive regulation of interleukin-2 production (Suppl. Table 6; Fig. 5D). We also observe more generic responses as evidenced by DEGs belonging to cell cycle checkpoints and tumor suppression annotated pathways (Fig. 5D).

## Discussion

Most avian disease studies have focused on specific immune cell types that support host response to a pathogen challenge ^19–24^. To the best of our knowledge, no studies have evaluated all cell types, a systems perspective, and no evaluations of avian single nucleus transcriptomes are available. A cell’s nucleus when cryopreserved adds a different analysis dimension compared to previous chicken single cell data with less signal associated with cellular stress, closer approximation of a cell’s status at the time of sampling, and enrichment for key transcription factors that regulate many downstream genes. Our primary goal was to first annotate all the major cell types, both immune and non-immune, found in the most predominant immune organ systems of bursa, thymus, and spleen, then use this newfound knowledge to examine an activation response in the evolutionarily novel avian bursal organ using a bacterial toxin to mimic systemic bacterial infections that occur in modern commercial poultry flocks.

The effects of infectious bursal disease virus (IBDV) infection on the bursa using scRNAseq data resulted in nine distinct clusters ^19^ that contrasts with our higher cluster diversity of 16 which can be explained by experimental differences in total sample size, treatment, nuclei versus live cells input, and likely other factors. The largest proportion of cells in both studies was B cells, as expected, but our stratification of subclustered B cell subtypes goes farther aided by utilzing overlap in cell and nuclei gene markers; some examples include *CD38, PTPRC,* and *XBP1* ^19^. Despite the informativeness of these shared bursal cell and nuclei markers substantial patterns of gene expression divergence exists. This again illustrates the subjective nature of cell type annotation and in some respects, the weakness of clustering techniques regardless of the targeted species. Fortunately, new thoughts and approaches are being discussed to move beyond traditional clustering ^52^. To better resolve B cell type developmental states we found subclustering to be necessary, but our annotation reliance on mouse B cell scRNAseq studies warrants caution in their future application to avian studies ^41,43^.

In all vertebrates, a well-functioning thymus is crucial for mounting effective immune responses against infections and diseases, yet detailed molecular profiles of all T cell types in the chicken are not available ^53^. The cellular composition of chicken thymus was first described in 1974 ^54^. Early thymocytes do not express CD4 or CD8 but naturally progress to the double positive (DP) stage expressing both *CD4* and *CD8A* until final commitment to differentiate into *CD4+* or *CD8+* T cells. We find these thymic stages of T cell development with the largest population being the DP T cells. Historically, chicken T cells have been identified at the protein level using antibodies to proteins, such as TCR2, which binds Vβ1 αβ T cells, yet with single cell or nuclei approaches, new hypothesis testing is possible. Our thymic nuclei transcriptome results, although just a preview showcase the complexity of T cell gene regulation. The addition of new immune gene nomenclature also promises to improve our discovery and curation of T cell subtypes ^55^. For instance, the chicken *FOXP3* gene, a marker for T reg cells, was fully characterized only recently and is highly expressed in secondary lymphatic organs but not in the thymus ^56^. When novel T cell molecular profiles in their varied states are coupled with gene knockout strategies, such as the deletion of αβ T cells showing a severe phenotype but not γδ T cells in chickens, the genetic sources of disease resistance can will become less vague ^57^.

Given the substantial economic losses to the poultry industry due to clinical and subclinical systemic bacterial infections, greater genetic insight regarding the functional roles of all cell types is imperative. How to define avian immune resilience to the thousands of bacterial species, some pathogenic, remains conceptually difficult to construct due to a need to model cell type sources of host immune tolerance ^58^. This first avian single nuclei study of host bursal response to a known bacterial toxin, LPS, establishes a preliminary accounting of the varied responses by cell type, immune and non-immune cell types. We find interesting novelties regarding the activated state of bursal B cell populations in the bursa and nominate certain ligand-receptor and signaling pathway candidates as modules of these complicated B cell networks when stressed with bacterial toxins. Our results reveal strikingly different interactions across different B cell subtypes but commonalities exist as well. In human, cellular communication inference evaluations importantly are shown to be largely concordant with alternative data modalities, such as intracellular signaling and spatial information ^59^ ^60^. In this study, the comparative inference of cellular crosstalk in the milieu of host immune response warrants caution but nonetheless offers a myriad of hypotheses to test in chickens experiencing bacterial infections ^61^. Of the many to consider, some of interest to us were interactions between *SEMA4D-CD72* and *PECAM1-CD38. SEMA4D* uses *CD72* as a functional receptor and enhances the activation of B cells by diminishing inhibitory signals from *CD72* that could be a consequence of LPS activation^62^. The *PECAM1-CD38* interaction may reflect a B cell activation to infection process since *CD38* has been postulated to act as a transmembrane receptor, whose interaction with its proposed ligand *PECAM-1* (*CD31*, a member of the Ig superfamily) results in proliferative and survival signals in lymphocytes ^63^. Building on earlier chicken cell-aggregate RNAseq studies in the bursa ^64^ ^65^, we now provide an preliminary interpretation of communicating B cell types when faced with a systemic bacterial infection.

In the bursa B cell, fate is driven by a careful synchrony of expressed genes such as *RAG*1 and *RAG2* both involved in V(D)J recombination that occurs in the embryonic spleen, and it is only “pre-B cells” with rearranged VDJ/VJ that home to the bursa, which is more an organ of expansion/selection than rearrangement ^66^. The V(D)J recombination activating genes, *RAG1* and *RAG2*, unlike in mammals, in chickens express high levels of the *RAG2* but not *RAG1* ^67^. The knockout of *RAG1* in chickens’ results in a lack of mature B and T cells ^68^. Our results offer complimentary evidence for cell type specific roles of *RAG1* and *RAG2*. Multiple formative B cell type themed hypotheses pertaining to their varied transcriptomic states within the evolutionarily novel avian bursal organ move us beyond antibody focused investigations in chicken. We chose the LPS challenge to mimic pathogenic bacterial infection because it is considered a well-established method to improve the characterization of innate immunity and readily comparable in future meta-analyses studies ^69^. Within this treatment narrative, many significantly expressed genes with cell-type specificity are of interest. MECP2 has some indications for a role in the inflammatory response ^70^; although not shown to be regulated in this study, we find genes affiliated with its signaling pathway that are such as MEF2C. Interestingly, MEF2C was found to play a role in LPS response, specifically regulating key B cell migration factors like PECAM1 and CXCR4 ^71^. CXCR4 is a critical chemokine receptor for B cell migration and maturation in the chicken bursa^72^.

*PAX5* gene expression, a critical transcription factor, is suggested to play a key role in controlling Salmonella infection in the commercial broiler, yet the key “hub” gene noted following enrichment testing was *FOXO3* ^73^. We do find *PAX5* expression diversity by cell type but not significant enrichment following LPS treatment. Other gene regulatory events perhaps of equal importance were noteworthy. *SYK* gene signaling in B cells, specifically the B pro BU1+ type, is an integral participant in the activation of Fcγ receptor-mediated phagocytosis in macrophages ^74^. SYK is a downstream signaling kinase for multiple Src family kinases including Lyn and Fyn in B cells, and Fcγ in macrophages - this is possibly an innate and adaptive responses integration role through its responsiveness to multiple upstream receptors in both innate and adaptive cell types. Given that SYK, is primarily expressed in mononuclear phagocytes and B cells, the finding in humans and mice that SYK gain-of-function variants are associated with immune deficiency put forward a hypothesis that should extensive natural sequence variation exist in the chicken models of selection could be developed ^75^.

In conclusion, this avian single-nuclei study expands our knowledge of immune cell types in the chicken bursa, spleen, and thymus, as well as their transcriptional responses to a bacterial toxin challenge in the novel bursa organ system. We identified gene expression signatures and their significant participation within immune-related pathways to provide new resources for future research in poultry immunogenetics and disease resistance. Furthermore, the evaluation of non-immune cell types gives us new perspectives on their often-overlooked role in communicating with various immune cell types, particularly those responsible for macrophage activation which is considered the most important first responder to bacterial infection. Understanding the intricate cell-to-cell and gene-to-gene interactions in the avian immune system is still a work in progress. These resources of the current study offer another point of understanding to move us closer to reaching a point of selecting on this variation to achieve true immune resistance.

## Limitations

Despite our novel findings for these organ systems, there is still a need for greater sampling depth by location, number, and genetic background, to name a few, but our annotation of chicken nuclei transcriptomes of major immune organ systems demonstrates this to be a rich resource for consistent immune cell type annotation and is consistent with a larger human study of immune cell types across diverse organs.

## Data and code Availability

The datasets generated during and/or analyzed during the current study are available in the Gene Expression Omnibus repository under accession GSE249652. The other data supporting this study are found within the article and supporting material.

## Acknowledgements

The authors thank the University of Missouri Genomics Technology Core for library preparation and sequencing. Computation for this work was performed on the high performance computing infrastructure provided by Research Computing Support Services and in part by the National Science Foundation under grant number CNS-1429294 at the University of Missouri, Columbia MO. Financial support was provided by the USDA NIFA Grant number 2020-67015-31574.

## Declarations and Funding Information

The authors declare there are no conflicts of interest. Financial support was provided by the USDA NIFA Grant number 2020-67015-31574.

## Author Contributions

W.C.W. and H.C. conceived and designed this study. C.H. did the animal work. E.R.S. and E.S.R. and S.K. performed computational analyses on the resulting data. E.R.S., C.H., J.S., J.K., A.B., S.J.L., H.C., Y.D., J.D., J.D., R.C.C., and W.C.W. annotated cell types. All authors participated in writing and editing the manuscript.

## Notes

### Competing Interest Statement

The authors have declared no competing interest.

