## Supplemental Figures for "A single-nucleus census of immune and non-immune cell types for the major immune organ systems of chicken"

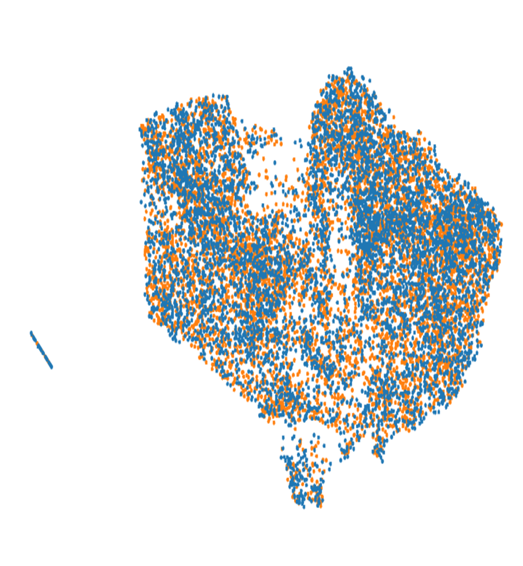

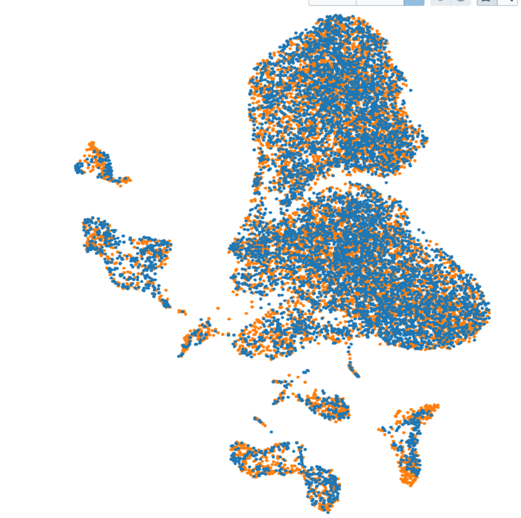

**Supplemental Figure 1.** Distribution of nuclei by treatment group in the bursa, blue is LPS and orange denotes control. A) all cell types B) subclustered B cells and spleen C) all cell types and D) subclustered lymphoid and myleiod immune cells identified in the initial clusters.

A)	B)

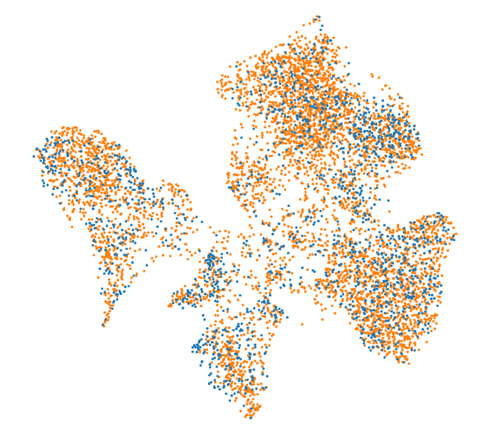

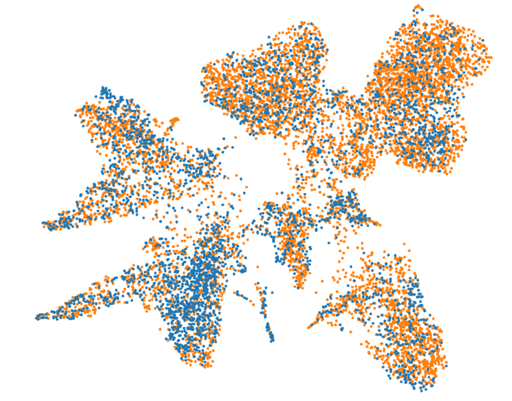
C)	D)

**Supplemental Figure 2.** Gene-specific expression plots across all cell types in the bursa. From left to right: *CD38, SOX4, ATXN1, CD72, PAX5, EBF1, RAG2, POLA1, TCF7, TP63, PPL, RHOV, KRT5, PDGFRA, THY1, CALD1, VWF, PTPRM, MYH11, MYLK, CACNA1C.*

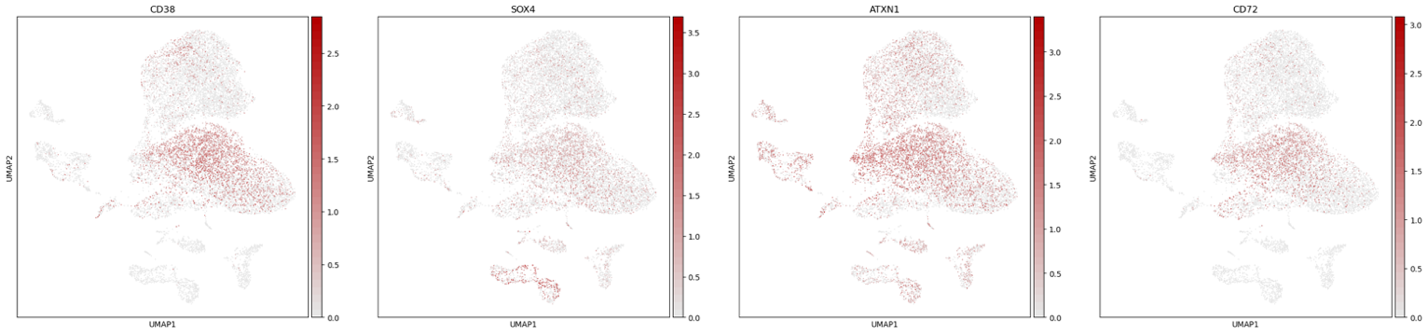

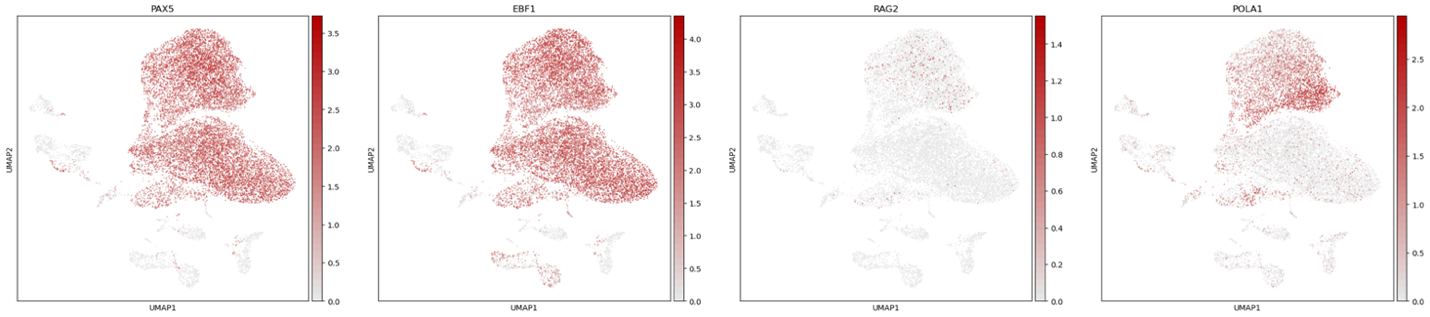

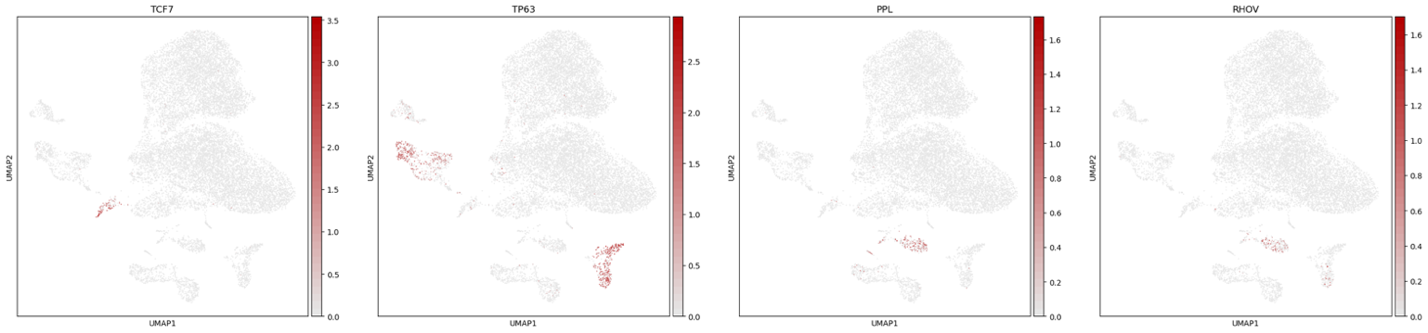

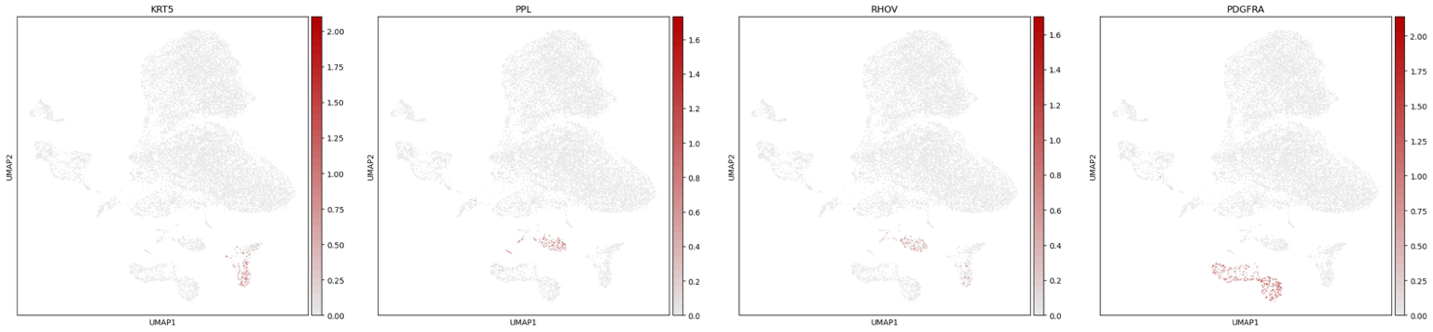

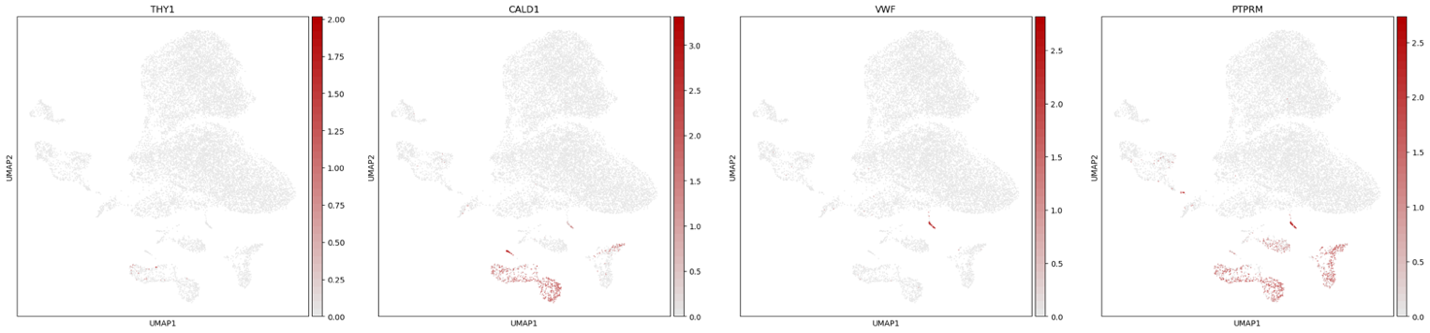

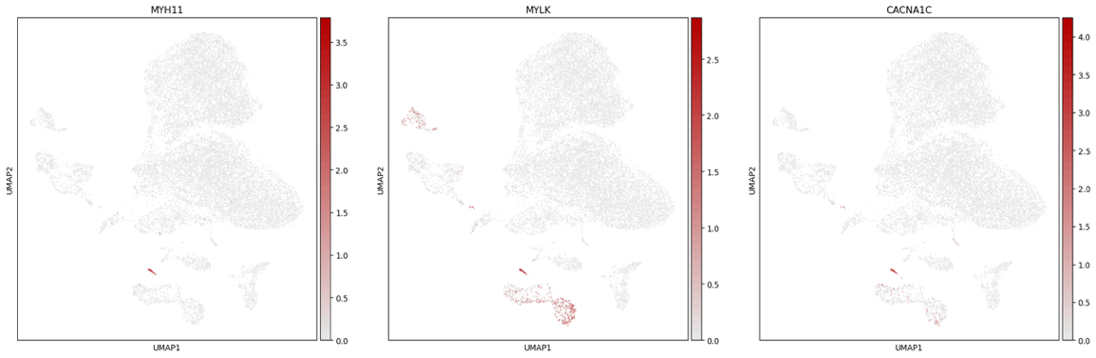

**Supplemental Figure 3.** Gene specific cell type annotation of only B-cell subtypes in the bursa. From left to right: *RAG2, TNFSF10, CD38, ATXN1, SOX4, CD72, POLA1, CXCR4, RUNX2, BCL2, EBF1.*

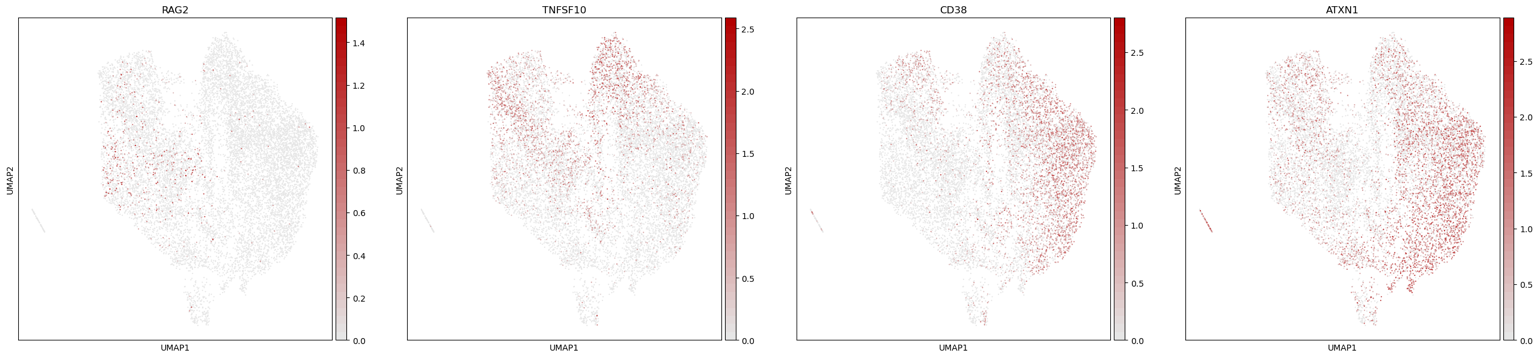

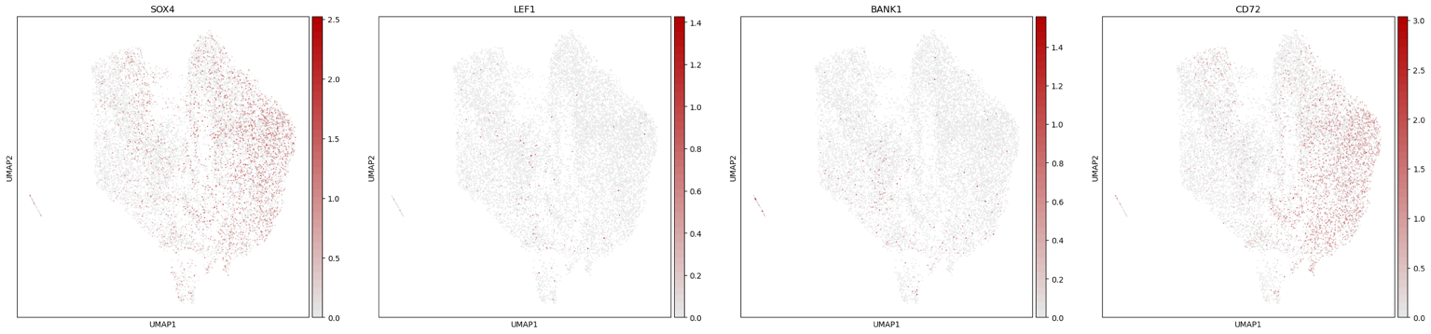

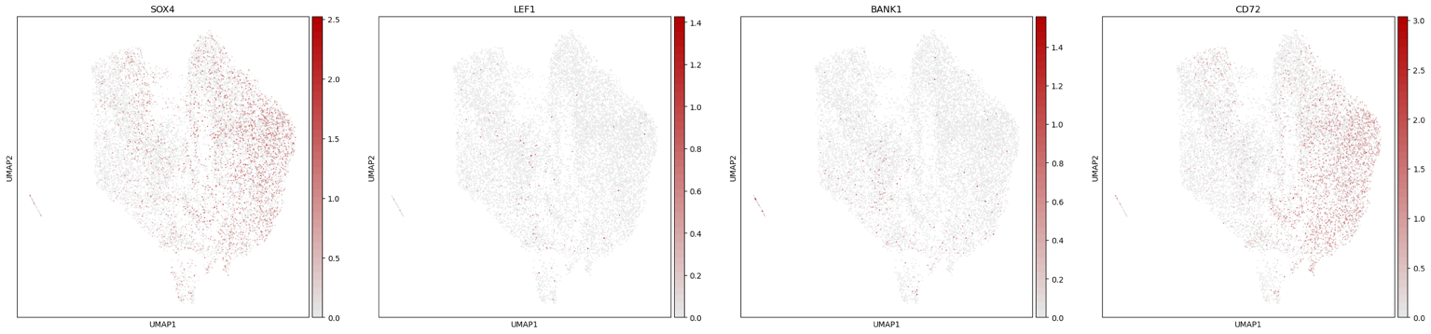

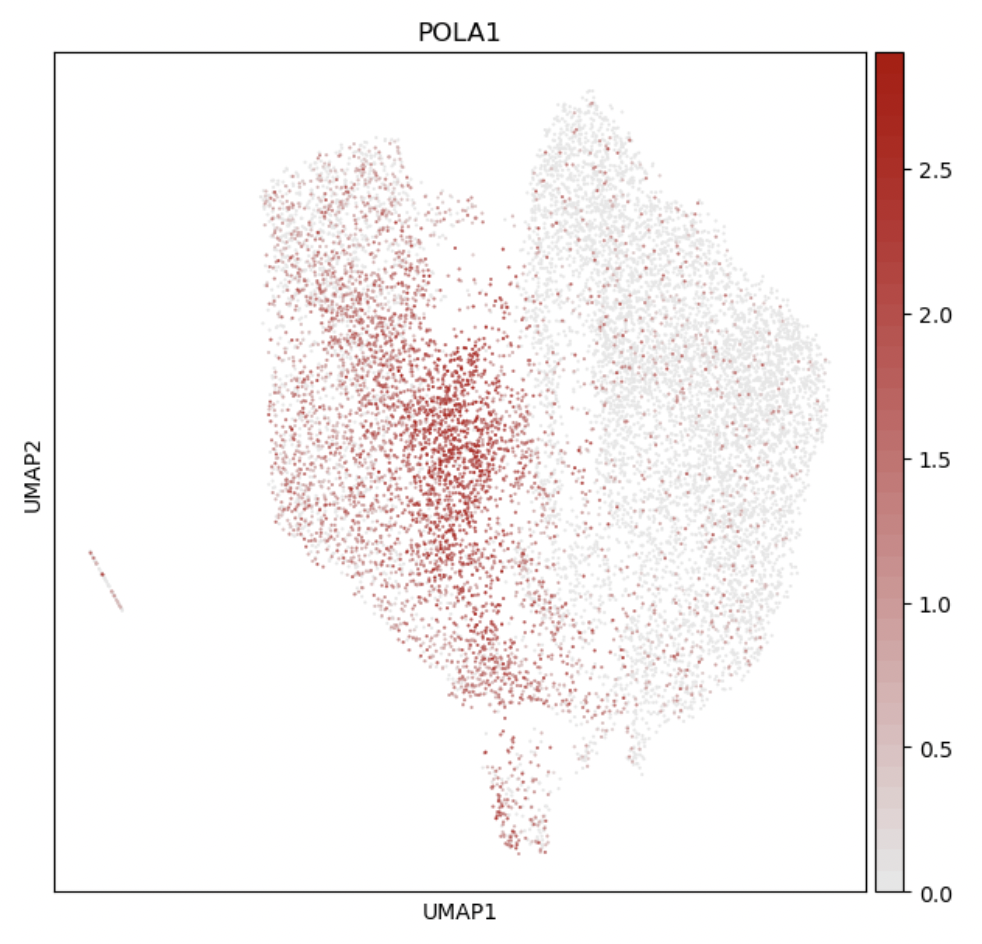

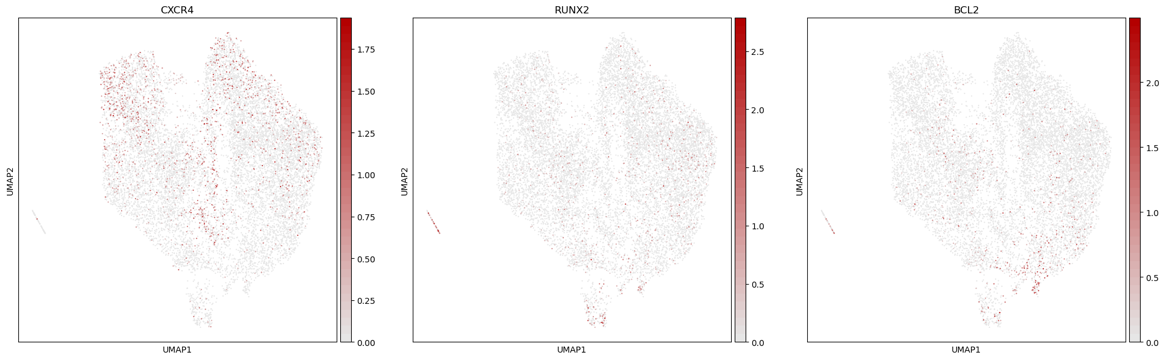

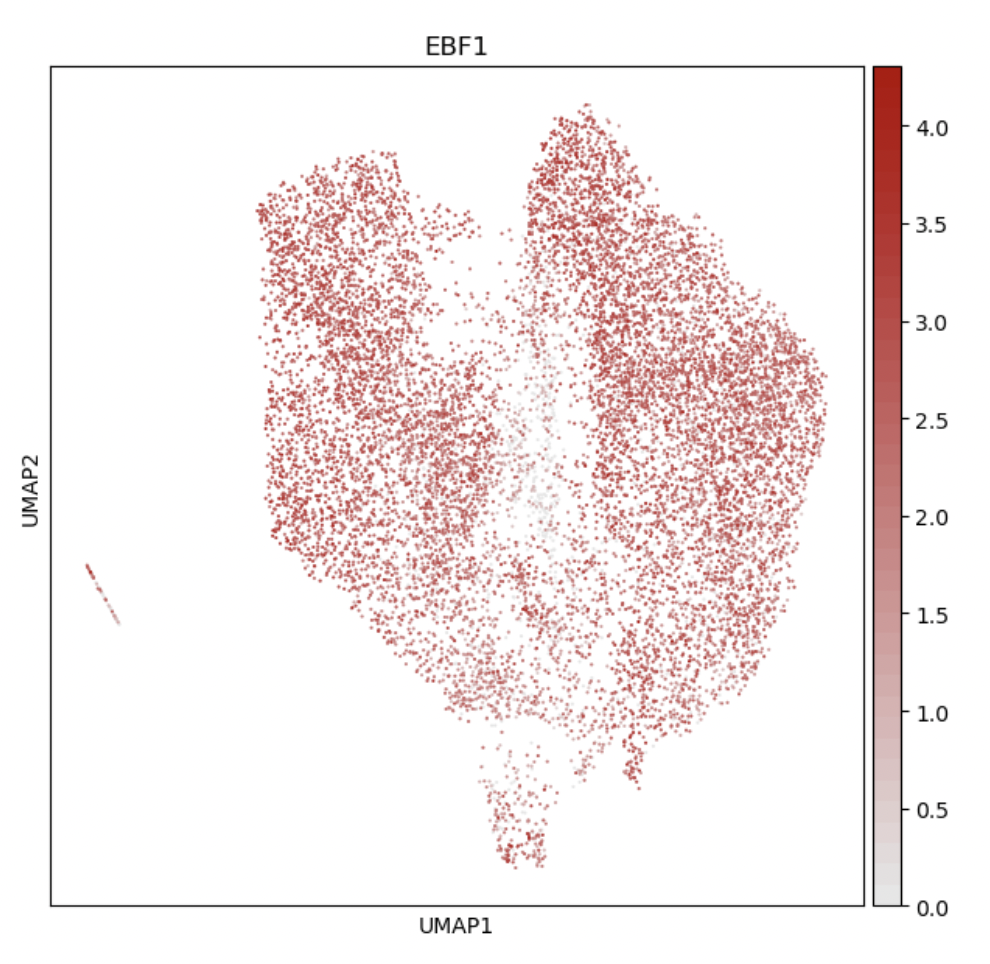

**Supplemental Figure 4.** Gene-specific expression plots across all cell types in the chicken spleen. From left to right: *CD3E, TARP, BCL11B, PAX5, BCL11A, EBF1, CD38, MARCO, CSF1R, CD86, FLT3, PDGFRA, VWF*.

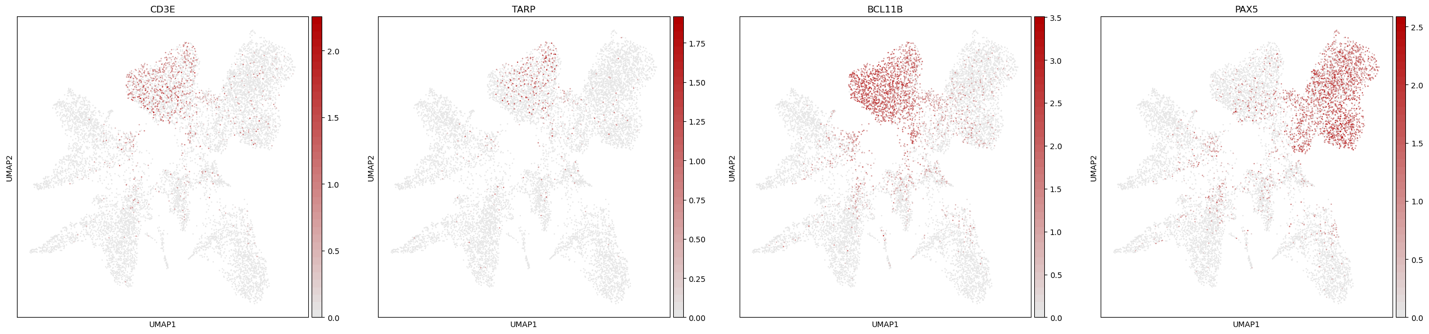

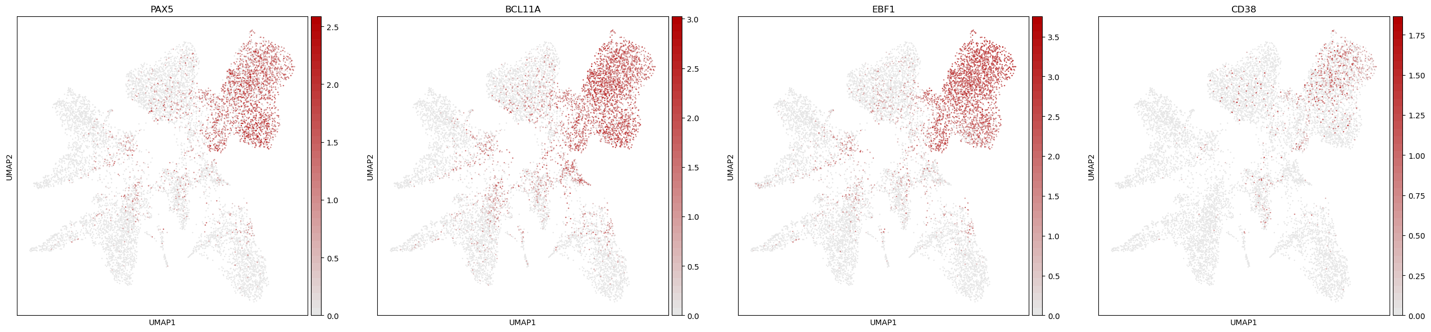

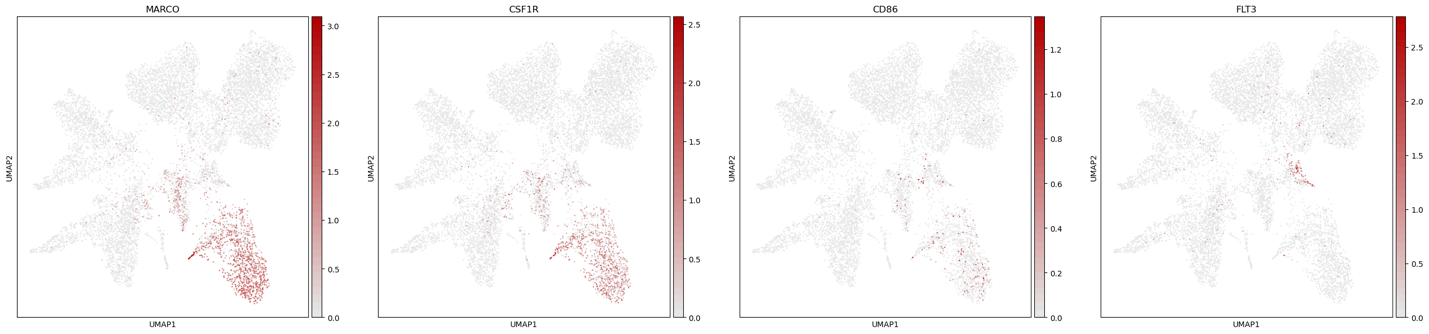

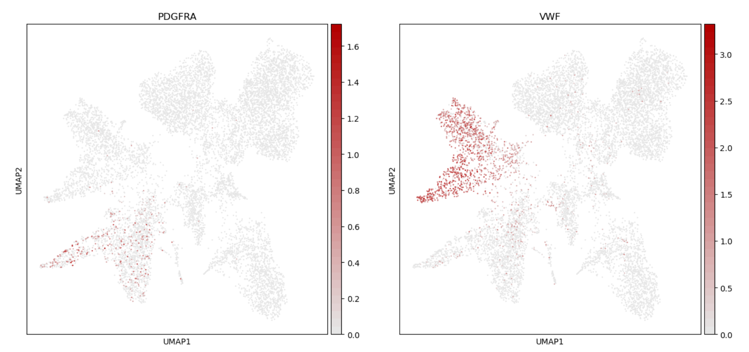

**Supplemental Figure 5.** Gene-specific expression plots across immune-only cell types in the chicken spleen. From left to right: *ENSGALG00010008265, ENSGALG00010003777, PAX5, RUNX2, ATXN1, EBF1, BCL11A, CD8A, CD4, GNLY, MARCO, CD86, CSF3R, TIMD4, MRC1, FLT3.*

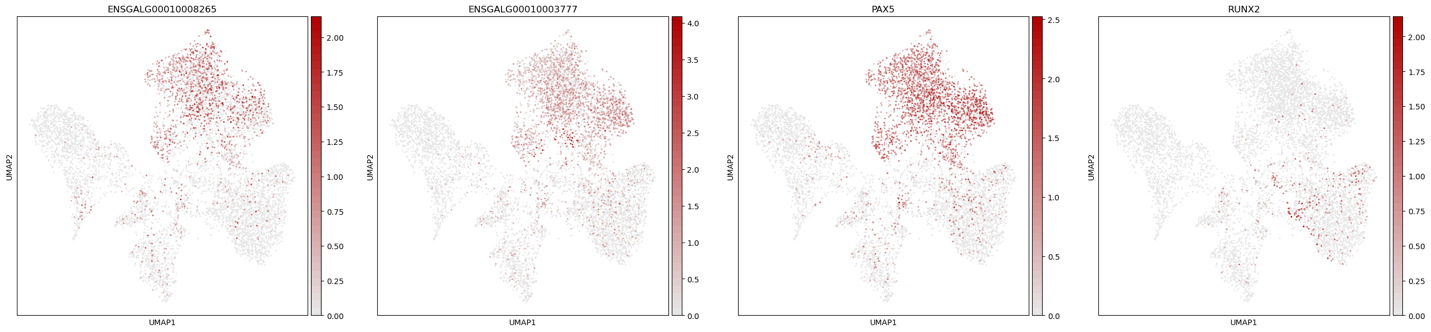

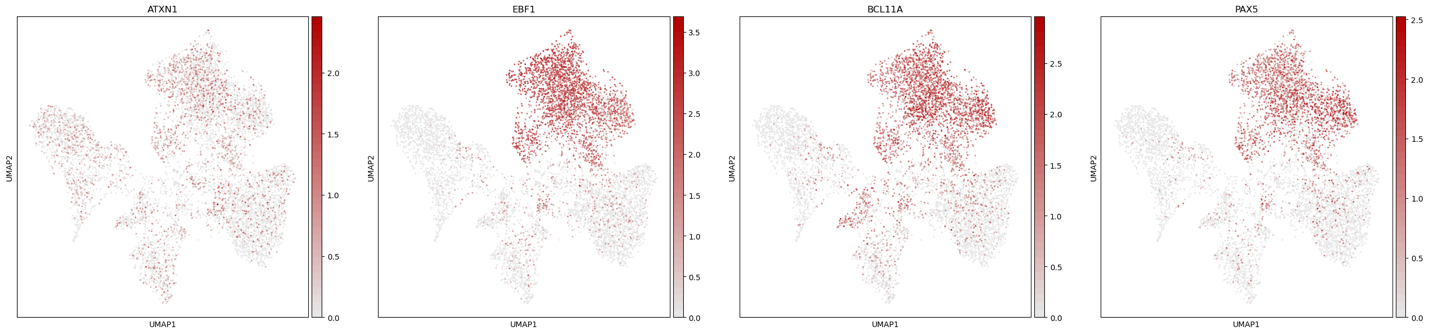

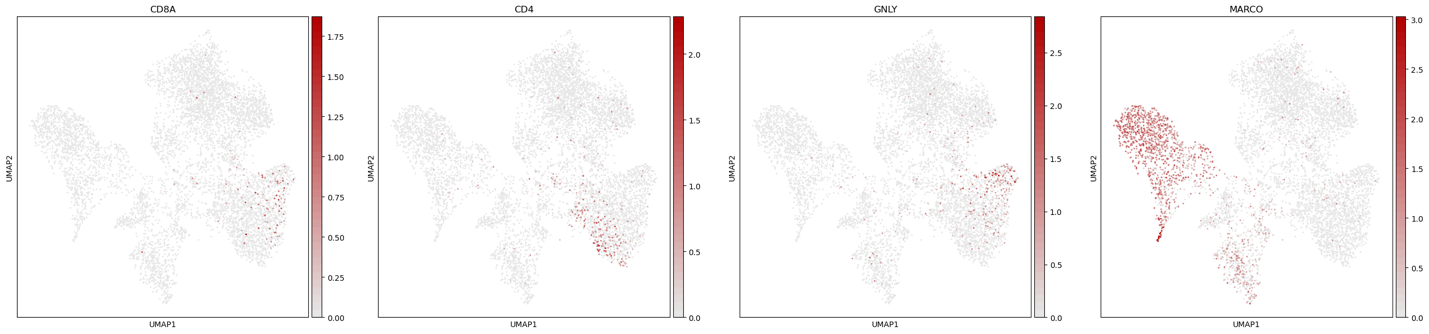

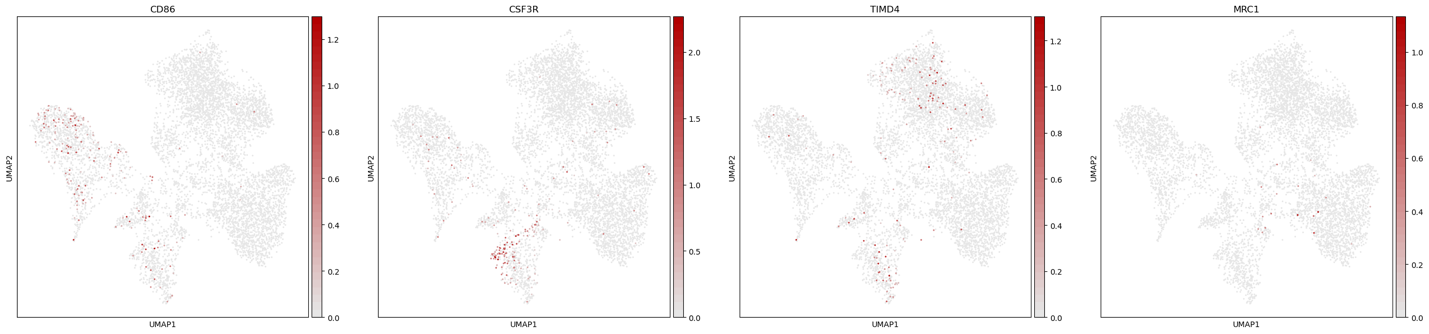

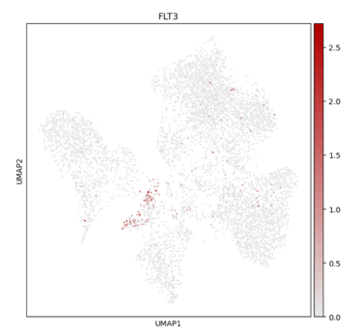

**Supplemental Figure 6.** Gene-specific expression plots across all cell types in the chicken thymus. From left to right: *TOX, POLA1, CENPF, ENSGALG00010025331 (TCR delta), TCRB, CTLA4, MYH9, IL7R, FLT3, BCL6.*

**Supplemental Figure 7.** Gene-specific expression plots across T cell types in the chicken thymus. From left to right: *CD4, CD8A, DNTT, CCR7, TCF7, BCL11B, TOX, POLA1, BRCA1, PLD5, MHCY11, TARP, CD274, CTLA4, TNFRSF9, LAG3, TNFRSF18, GATA3.*

**Supplemental Figure 8.** Gene expression overlap by the largest B cell clusters. List 1 is pre pro B, list 2 is proliferating B, list 3 is B cells (1) and list 4 is large pro-B cells as identified in all cell types clusters in Fig. 2A.

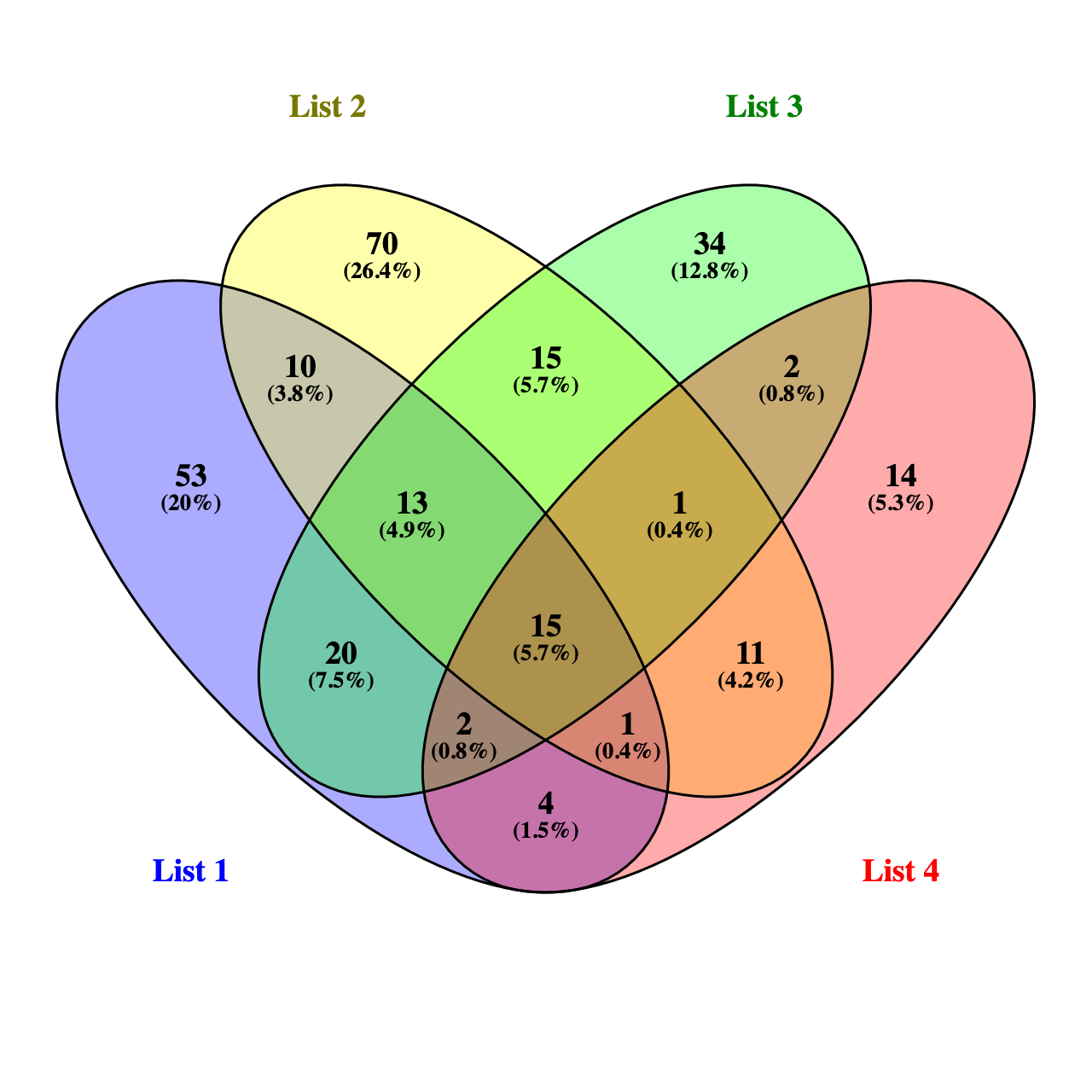
